# Transcription start site heterogeneity and its role in RNA fate determination distinguish HIV-1 from other retroviruses and are mediated by core promoter elements

**DOI:** 10.1101/2023.05.22.541776

**Authors:** Siarhei Kharytonchyk, Cleo Burnett, Keshav GC, Alice Telesnitsky

## Abstract

HIV-1 uses heterogeneous transcription start sites (TSSs) to generate two RNA 5’ isoforms that adopt radically different structures and perform distinct replication functions. Although these RNAs differ in length by only two bases, exclusively the shorter RNA is encapsidated while the longer RNA is excluded from virions and provides intracellular functions. The current study examined TSS usage and packaging selectivity for a broad range of retroviruses and found that heterogenous TSS usage was a conserved feature of all tested HIV-1 strains, but all other retroviruses examined displayed unique TSSs. Phylogenetic csomparisons and chimeric viruses’ properties provided evidence that this mechanism of RNA fate determination was an innovation of the HIV-1 lineage, with determinants mapping to core promoter elements. Fine-tuning differences between HIV-1 and HIV-2, which uses a unique TSS, implicated purine residue positioning plus a specific TSS-adjacent dinucleotide in specifying multiplicity of TSS usage. Based on these findings, HIV-1 expression constructs were generated that differed from the parental strain by only two point mutations yet each expressed only one of HIV-1’s two RNAs. Replication defects of the variant with only the presumptive founder TSS were less severe than those for the virus with only the secondary start site.

## Introduction

HIV-1 proviruses contain a single transcriptional promoter in their upstream long terminal repeat (LTR). This single transcription unit generates RNAs that are either spliced into one of HIV-1’s multiple mRNA species or else transported unspliced from the nucleus to the cytoplasm. Unspliced HIV-1 RNAs serve either as mRNAs for virion structural proteins or as the genomic RNAs packaged into virions.

HIV-1 primary transcripts were long regarded as a single RNA species, but it was more recently found that they exist as two distinct 5’ isoforms in cells, only one of which (^cap^1G) is encapsidated. The other (^cap^3G) is retained in cells and serves intracellular functions [1, 2]. This multiplicity of RNA forms is achieved through the usage of clustered transcription start sites (TSS) located at the border of the U3 and R regions of the viral LTR. The two RNAs begin with either one or three guanosines, depending on which of three sequential Gs in the GGG motif at the U3/R border is chosen to initiate transcription. Despite differing by only two nucleotides, these RNAs perform highly divergent functions. Only the ^cap^1G form is packaged into nascent virions, while ^cap^3G RNAs are enriched on polysomes and among spliced RNAs [1, 3].

NMR work reveals that these functional differences reflect different RNA structures [4]. RNA fragments’ abilities to dimerize differ for RNAs with 1 or 3 Gs at their capped 5’ ends. ^cap^1G RNAs feature an exposed DIS stem-loop that allows the “kissing” interactions that initiate RNA dimerization with a second ^cap^1G RNA, and structure that unmasks nucleocapsid protein (NCp) binding sites [1, 5, 6]. The DIS loop is sequestered by intramolecular basepairing in the ^cap^3G conformer, which also differs from ^cap^1G RNA in its exposure of the major 5’ splice site for spliceosome recruitment. Additionally, unlike ^cap^1G RNAs, where RNA structure renders cap moieties inaccessible to cap binding factors, the extended 5’ ends of ^cap^3G RNAs are accessible for translation initiation [6].

The pivotal role TSS choice plays in dictating HIV-1 RNA fates adds to the importance of transcriptional initiation as a key HIV-1 regulatory step. HIV-1 transcription is regulated by an enhancer/promotor region in the LTR’s U3 region and is driven by the host’s RNA polymerase II (Pol II) machinery. When classified by TSS use patterns, there are two main types of Pol II promoters: focused and dispersed [7, 8]. A dispersed promoter contains multiple TSS dispersed over a 50 to 100 nucleotide region, while a focused core promoter contains a single predominant TSS. A variation of this latter class is twin-TSS promoters with two well defined TSS spaced by 0-3 bp [9].

Although the mechanisms are still debated, it is believed that core promoter element binding factors phase Pol II and determine TSS choice (reviewed in [10, 11]). The TATA-box is one candidate element for modulating focused TSS choice [7, 9]. Canonical TATA-boxes have a variable TATAWAAR consensus [12, 13] with many Pol II promoters completely lacking discernable TATA-boxes [14, 15]. Once the TATA-box-binding protein (TBP) of transcription factor IID (TFIID) binds the core promoter, a cascade of other factors plus Pol II are recruited, ultimately leading to transcriptional preinitiation complex formation [11]. Some focused promoters with TATA-boxes also have a motif called the Initiator (Inr) [16]. Inr flanks the TSS, is recognized by the TAF1 and TAF2 subunits of TFIID, and plays roles in transcription activation, direction, and TSS determination [12, 16]

It was long believed that retroviruses used focused promoters with single TSSs. This was based on outcomes of reverse transcription, which require genomic RNA 5’ ends to match the RNA’s 3’ end. Although homology between RNA 5’ and 3’ ends alone is sufficient to guide reverse transcriptase strong stop template switching, the precise sequences and secondary structures of viral RNA termini greatly facilitate this process, which occurs at a single template position for both HIV-1 and simple retroviruses [17–20]. Although the discovery that HIV-1 generates two RNA 5’ isoforms at first seemed to challenge the need for precise ends, the packaging of only one RNA species provides a mechanism for maintaining the precise homology required for reverse transcription fidelity.

Two HIV-1 isolates, NL4-3 and MAL, have been shown to display TSS usage heterogeneity [1, 5]. Both are HIV-1 group M members, with NL4-3 being subtype B and MAL a recombinant with subtype A 5’ leader sequences. However, these strains represent a very small portion of HIV-1 diversity. Group M is subdivided into subtypes A through L and more divergent HIV-1 groups (N, O, and P), each from an independent zoonotic transmission, also have been described [21, 22]. Here, we examined the prevalence of heterogeneous TSS usage among retroviruses, defined viral features that specify heterogeneous TSS usage, and used this knowledge to express unique HIV-1 RNA isoforms and their functional consequences.

## Results

### Heterogeneous TSS usage differentiates HIV-1 strains from other retroviruses

In the two HIV-1 strains tested thus far for heterogenous start site use - one each from subtypes B and A - their TSS regions possess a trinucleotide motif consisting of three sequential guanosines, with the first and third recognized to initiate transcripts alternately with three or one 5’ terminal guanosine [1, 5]. Thus, the conservation of this motif was tested a first step toward addressing TSS usage among lentiviruses.

TSS region sequences were analyzed for more than 3000 near full-length LTR sequences available in the Los Alamos HIV-1 Sequences Database (https://www.hiv.lanl.gov/content/index). Analyzed sequences included representatives of HIV-1 groups M, O, N, and P, plus several isolates of SIVcpz and SIVgor. The results of this analysis **(Fig.1A)** revealed that a large majority (approx. 99%) retained three sequential guanosines, flanked by additional conserved residues to yield a consensus sequence of TACT **GGG** TCTC. Thus, HIV-1’s TSS region is highly conserved.

**Figure 1.**
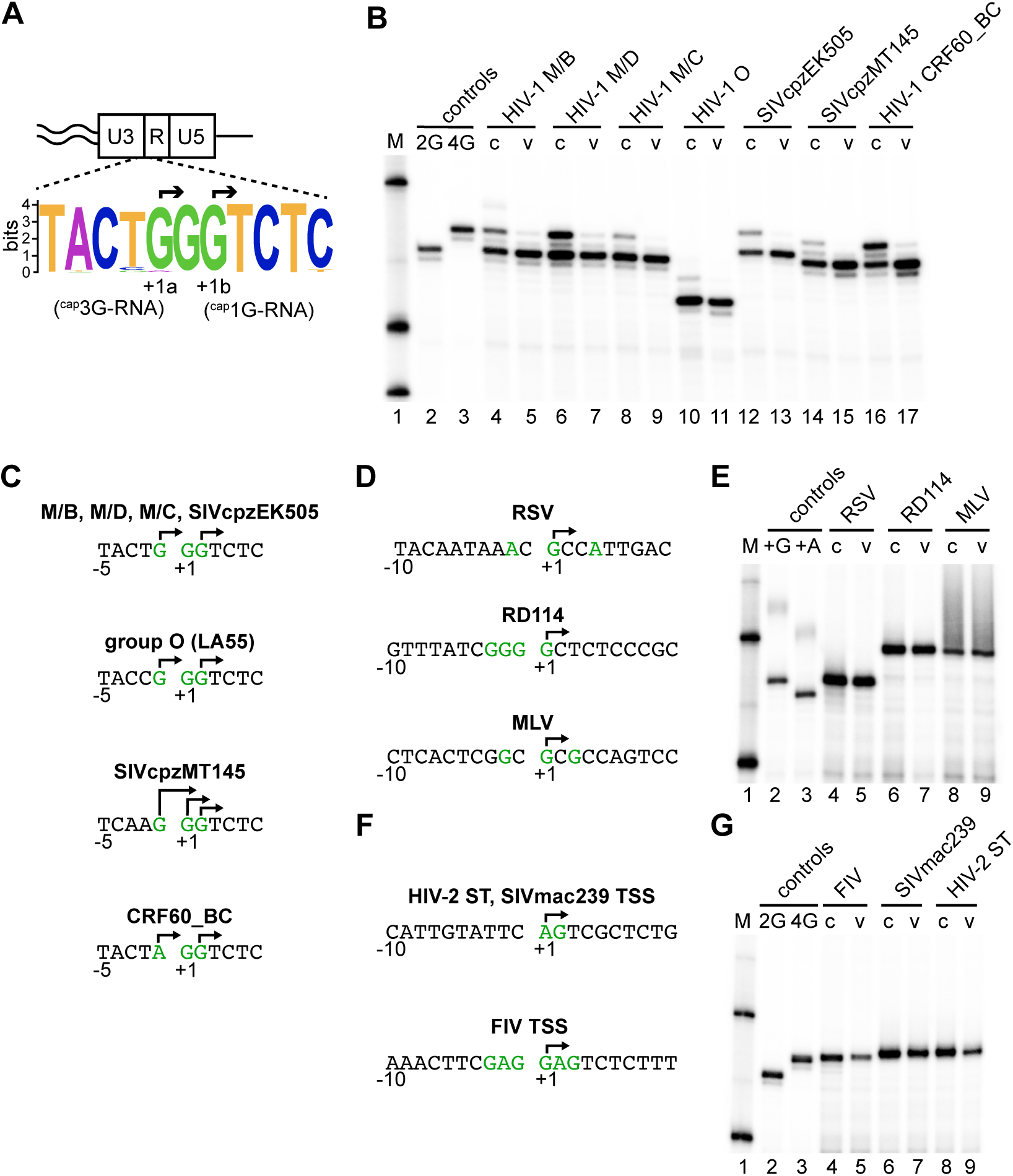
Extents of heterogeneous TSS usage among retroviruses and a comparison of cellular and virion RNA 5’ ends. (A) Schematic representation of conserved sequences at the HIV-1 U3/R border. Consensus was obtained from aligning 575 isolates’ 5’ LTR sequences from the Los Alamos HIV-1 Sequences Database, using WebLogo software. Overall height of each stack indicates the sequence conservation at that position measured in bits, whereas the height of symbols within the stack reflects the relative frequency of the corresponding base at that position [45]. Note that HXB2 (accession # K03455.1) GenBank coordinates indicate HIV-1 mRNA starts at the second G of the GGG motif, at LTR position 455, but more recent studies indicate 455 is the least commonly used TSS in the GGG motif [1, 2]. Thus, to preserve HXB2 numbering while reflecting experimentally validated start sites, +1 was used to indicate the beginning of viral RNA according to GenBank and the two major experimentally determined TSSs are labeled +1a and +1b. (B) Single base resolution assay results of RNA 5’ ends measured by CaDAL. M, size marker; lanes 1 and 2, control PCR products from 2G and 4G LTR plasmids, which co-migrate with CaDAL products of ^cap^1G and ^cap^3G RNAs, respectively; lanes 3-16, RNA samples extracted from transfected cells (c) or virus media (v). HIV-1 groups, subtypes or strains are indicated: specific isolates are in Materials and Methods. Note that the smaller size of group O CaDAL products reflects primer design. (C) TSS for different HIV-1 isolates, confirmed by sequencing of CaDAL products, indicated by arrows. (D) TSS usage for simple retroviruses and non- HIV-1 lentiviruses. Sequences surrounding the TSSs of RSV, RD 114 and MLV. Purines located in the TSS vicinity are represented in green. Experimentally determined TSSs are indicated with arrows. (E) Single base resolution assay of the simple retroviruses’ RNA 5’ ends, as assessed by cap-dependent adaptor ligation/PCR. M, molecular size marker; lanes 1 and 2, control PCR products, lanes 3-8, 5’ end products of RNA samples extracted from chronically infected cells (c) or virus media (v). Samples containing RSV, RD114 or MLV RNA are indicated. (F) Sequences surrounding the TSSs of HIV-2 ST, SIVmac239 and FIV. Purines that potentially could be used as TSSs are shown in green. Experimentally determined TSSs are indicated with arrows. (G) Single base resolution assay of lentiviruses’ RNA 5’ ends assessed by CaDAL. M, molecular size marker; lanes 1 and 2, PCR products for 2G and 4G HIV-1 control plasmids; lanes 3-8, products from RNA samples from cells (c) or virus media (v). Samples containing FIV, SIVmac239 or HIV-2 ST RNA are indicated.

To address if this conservation correlated with heterogeneous TSS usage and RNA fate determination, intracellular transcripts and virion RNAs generated by six viruses, selected to represent the breadth of HIV-1 strains and their close relatives, were compared to those generated by NL4-3 (**Fig. 1B)**. RNA 5’ ends were determined with single base precision for the intracellular transcripts and virion RNAs generated by subtypes C and D isolates; by a group O isolate; by SIVcpzEK505, which is closely related to HIV-1 group N [23]; by SIVcpzMT145, which is not associated with any known human HIV-1 infections [23], and by a circulating recombinant form (CRF60_BC) selected due to its rare TSS motif (AGG in place of GGG). For five of these, approximately 1 kb of strain-specific sequences, including the 5’ LTR through initial sequences in *gag,* replaced the corresponding sequences of an NL4-3 -based vector. These vectors were mobilized by co-transfection with a NL4-3 based λϑΙ- helper. The subtype C isolate was tested using an infectious clone.

A **ca**p-**d**ependent **a**daptor **l**igation/PCR protocol (CaDAL assay) [3, 24] was used to map precise RNA 5’ ends. This assay was designed to exclude 5’ truncated products from subsequent amplification, thus minimizing contributions from partially degraded RNAs, prematurely terminated cDNAs, or cDNA extended by the terminal deoxynucleotidyl transferase-like activity of reverse transcriptase. This technique involves first generating cDNA to viral RNA, then ligation to an adaptor that relies on the RNA 5’ cap of the resulting cDNA/RNA duplex. Primers used for first strand DNA synthesis annealed downstream of HIV-1’s 5’ splice site. As a result, the assay reported capped 5’ ends of unspliced transcripts only.

Assay results **(Fig. 1B;** c indicates cell lanes) showed that all tested HIV-1s produced two prominent intracellular RNAs, while SIVcpz MT145 produced three 5’ isoforms. Sequencing CaDAL products (**Fig. 1C**) showed that the two intracellular RNA forms for most strains were ^cap^1G and ^cap^3G, with CRF60_BC RNAs bearing ^cap^1G and ^cap^AGG ends, as expected due to its AGG TSS motif. The three RNAs produced by SIVcpz MT145 had ^cap^1G, ^cap^2G, and ^cap^3G ends. Although all these HIV-1-type viruses displayed heterogeneous TSS usage, RNA forms’ ratios varied, with group O producing the highest and CRF60_BC the lowest relative amounts of ^cap^1G RNAs. In data not shown, properties of a CRF60_BC variant with a GGG motif and of an NL4-3 variant with an AGG motif suggested the rare A in the CRF60_BC AGG motif was responsible for its skewed TSS ratios.

Despite their multiple intracellular RNA isoforms, all or nearly all encapsidated RNAs observed for these viruses possessed ^cap^1G 5’ ends, as had previously been reported for subtypes A and B RNA packaging (**Fig. 1B;** v indicates viral RNA lanes). Thus, both heterogeneous TSS usage and regulation of viral RNA fates by alternate 5’ ends are conserved among HIV-1 strains.

To address the conservation of these properties among more distally related retroviruses, TSS usage was determined for the gamma-retroviruses Moloney murine leukemia virus (MLV) and RD114 (a feline endogenous retrovirus [25]) and for the alpha-retrovirus Rous sarcoma virus (RSV). MLV and RSV have been studied for decades and older analyses of their virion RNAs suggest that both use unique TSSs [26, 27]. However, there is no data on RNA 5’ ends in infected cells, and both RSV and MLV contain several purines near their defined TSSs that could serve as alternate starts **(Fig. 1D)**. RD114 has a run of four Gs near its putative TSS: a property it shares with the gamma-retrovirus Spleen Necrosis Virus and with the Visna-Maedi lentivirus. Although its TSS has not been mapped to our knowledge, RD114 was selected for study based on its potential to produce multiple RNA forms **(Fig. 1D)**.

To generate infected cell and virion RNA samples for these additional viruses, virus-specific host cell lines were chronically infected with RSV, RD114 or MLV viruses (see Material and Methods), and CaDAL was used to determine precise RNA 5’ ends **(Fig. 1E)**. The results showed that all three of these simple retroviruses produced single RNA forms within cells, and that these same RNAs also were packaged into virions. Sequencing CaDAL products confirmed that both RSV and MLV transcripts initiated at the G residues that had previously been implicated [26, 27] **(Fig. 1D)**. RD114 was determined to initiate RNA synthesis uniquely from the final G residue of the 4 G run at its TSS **(Fig. 1D)**.

After finding that simple retroviruses differed from HIV-1 in TSS usage and packaging, and thus apparently in their strategies for defining genomic vs messenger RNA pools, heterogeneous TSS usage was then examined for additional lentiviruses. These included two viruses belonging to the HIV-2/SIVmac clade (HIV-2 ST and SIVmac239) plus the feline lentivirus FIV. As indicated in **Figure 1F**, TSS region sequences for these viruses differ from those conserved among HIV-1 strains but retain significant purine richness.

To map the 5’ ends of these lentiviral RNAs, SIVmac239 and FIV cell and viral RNAs were harvested from chronically infected human and cat cell lines, respectively, while HIV-2 was transiently expressed in 293T cells (see Materials and Methods). RNA extracted from producer cells and viral particles was subjected to CaDAL **(Fig. 1G)**. The results showed that a single RNA 5’ isoform was present in cells for each lentivirus, and the same RNA was encapsidated **(Fig. 1G)**. Sequencing CaDAL products confirmed that all detectable transcription initiated from the first of two TSS-region purines for both HIV-2 and SIVmac239, yielding an RNA initiating in a 5’ A, and that FIV transcription initiated exclusively from the fourth nucleotide (G) in the six-purine run at its TSS **(Fig. 1F)**. Thus, neither heterogeneous TSS usage nor the use of distinct RNA species in packaging was observed for these non-HIV-1 lentiviruses.

Maximum-likelihood phylogenetic analysis was then conducted to compare observed patterns of TSS usage to the phylogenetic relatedness of these viruses, using sequences spanning from the TATA element of the core promoter (positions -30 for HIV-1 and FIV, and -31 for HIV-2 **(Fig. 2A)**) through the *gag* start codon. The resulting phylogenetic tree was similar to previously published relationships for primate lentiviruses based on *gag* and *pol* sequences **(Fig. 2B)** [28], with all HIV-1 strains clustering together, a separate branch for the HIV-2 viruses, and FIV the most distantly related of the compared lentiviruses. Interestingly, the grouping of viruses on the phylogenetic tree coincides with heterogeneous *vs* unique TSS usage **(Fig. 2B)**. This correlation suggested the possibility that the strategy of regulating RNA fates through heterogeneous TSS usage emerged after separation of the SIVcpz/HIV-1 and SIVsm/HIV-2 lineages [28] and before radiation of the SIVcpz/HIV-1 lineage into existing HIV-1 and HIV-1-related viruses.

**Figure 2.**
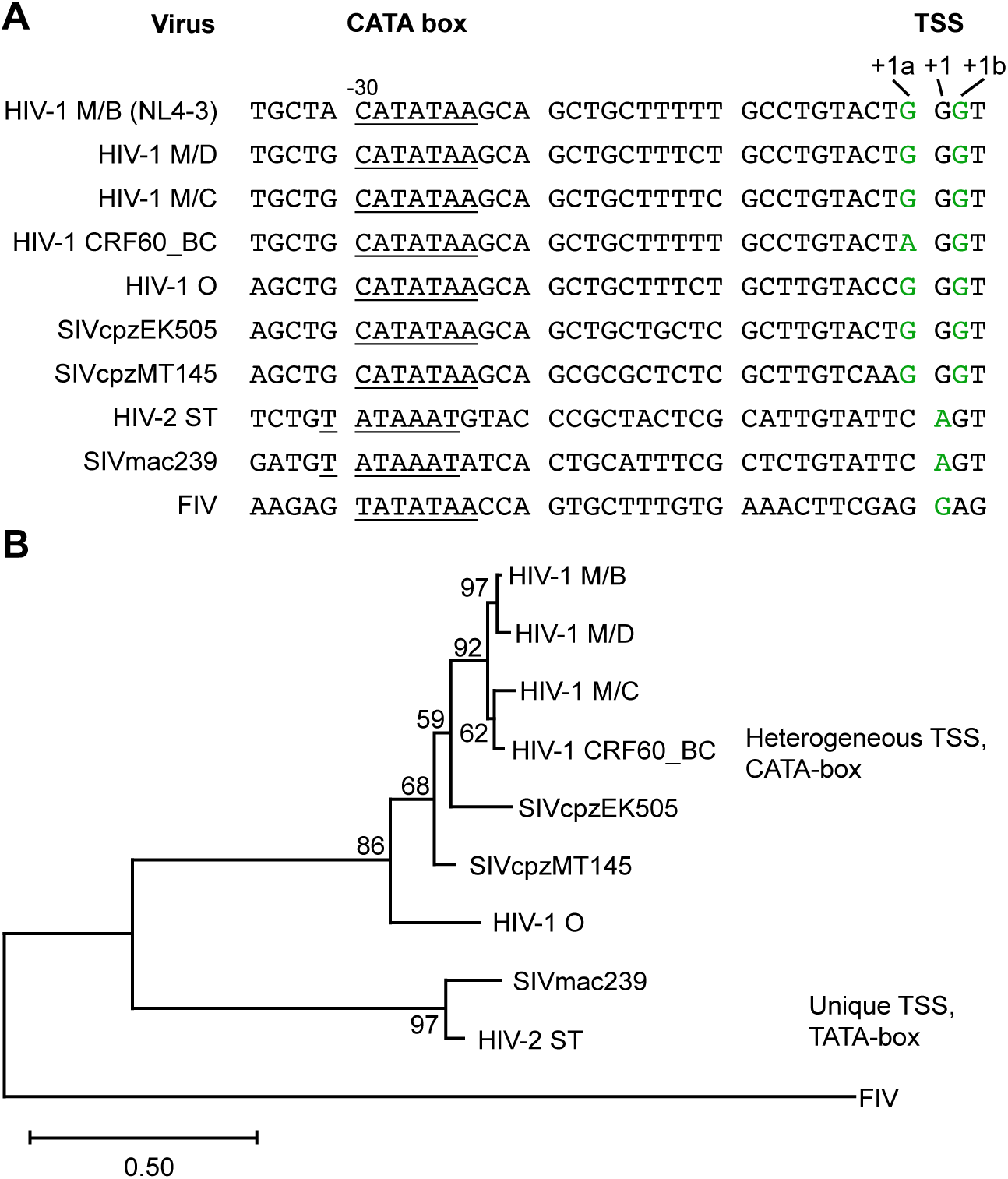
Phylogenetic comparison of core promoter elements in lentiviruses. Correlations of these viruses’ TATA elements, TSS usage and phylogenetic relationships. (A) Sequences from -35 to +3 of the lentiviruses included in this study. CATA or TATA box motifs are underlined and experimentally determined TSSs are indicated in green. (B) Maximum likelihood phylogenetic tree depicting the relationship of the tested lentiviruses based on core promoter and 5’ untranslated region sequences. The tree was built using the General Time Reversible model [47] with gamma distribution. FIV was used as the outgroup. Numbers next to the nodes represent branch support (1,000 bootstrap replicates).

### Major determinants of TSS heterogeneity are located upstream of the TSS

Previous work demonstrated that altering HIV-1 TSS-region G stretch lengths shifts start site positions but maintains heterogeneous TSS usage, suggesting that start site choice is at least partially based on distance counting from upstream promoter elements [1]. These findings suggested that the major determinants of TSS heterogeneity were located upstream of the TSS. Thus, with knowledge that retroviruses other than HIV-1 displayed unique TSS usage, LTR chimeras were generated **(Fig. 3A)** to test if HIV-1 sequences upstream of the TSS were sufficient to confer heterogeneous TSS usage.

**Figure 3.**
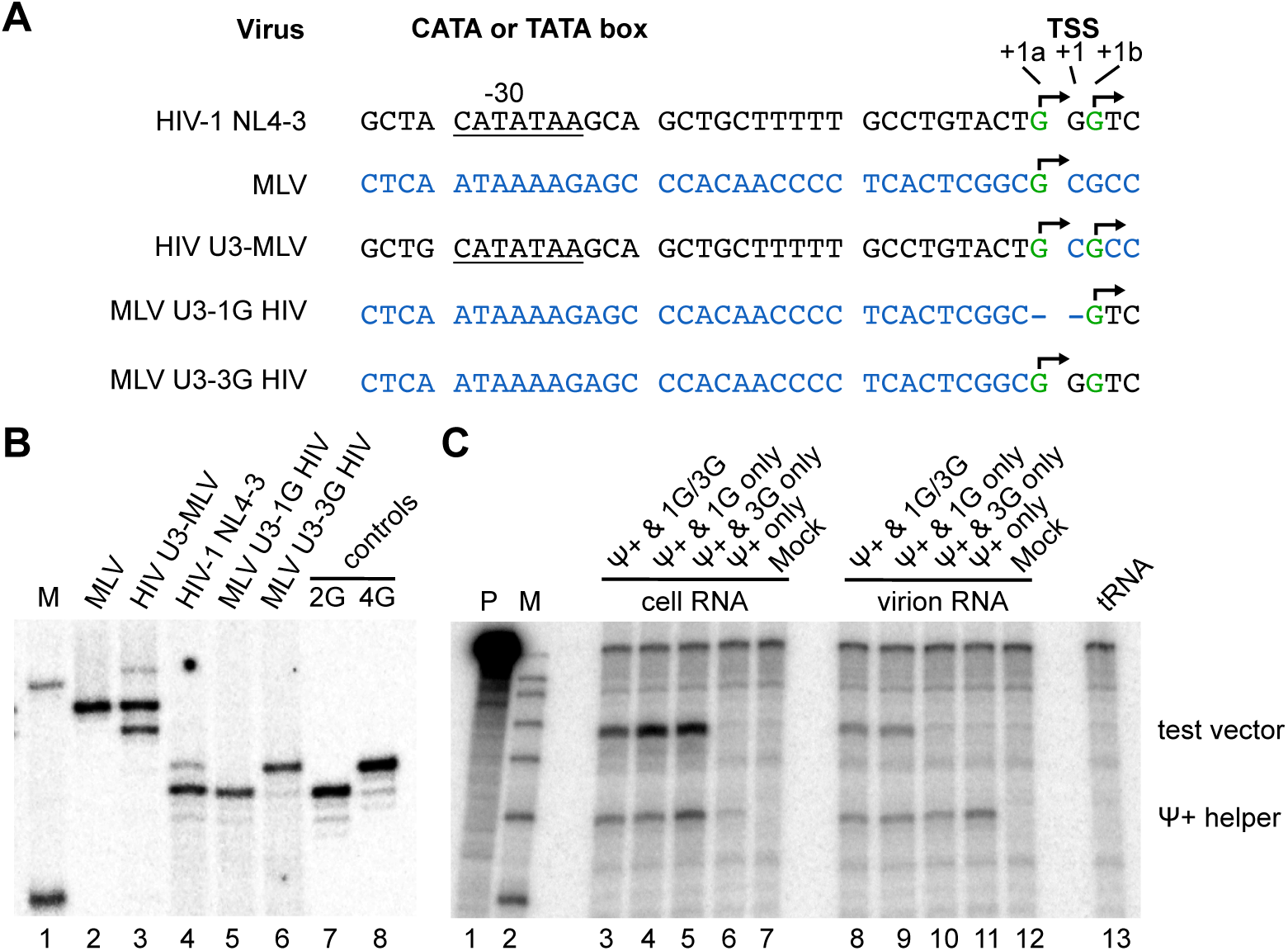
U3 exchange chimeras to test sequences upstream of TSS for roles in heterogeneous TSS usage. (A) Sequence of the MLV and HIV-1 vectors with chimeric promotors under control of HIV-1 U3. HIV-1 sequence shown in black, MLV sequence shown in blue. Observed TSS shown in green and indicated with arrows; (B). Single base resolution assay of the 5’ ends of the viral RNA produced in the cells transfected with HIV-1 or MLV vectors and assessed by cap-dependent adaptor ligation/PCR. M, molecular size marker; lines 2 and 3, PCR products for 2G and 4G HIV-1 control plasmids; line 4, RNA sample extracted from cells transfected with HIV-1 NL-4-3 vector under HIV-1 U3; lines 5-6, RNA samples extracted from cells transfected with chimeric HIV-1 vectors under control of MLV promoter with HIV-1 sequence begins with one (line 5) or three Gs (line 6); lines 7 and 8, RNA sample extracted from cells transfected with MLV vectors, under MLV or HIV-1 promoters correspondingly (C) Cellular expression and packaging of MLV U3 driven HIV-1 vectors RNA in competition with Ψ+ helper as monitored by RNase protection assay. RNA samples obtained from transiently transfected 293T cells (lanes 3-7) or extracted from virus containing media (lanes 8-12) were hybridized with chimeric riboprobe protecting 201nt fragment of the helper and 289 nt fragment of the test vectors (indicated on the right). Lane designations indicate which vectors were co-transfected with helper or helper transfected alone; mock: mock transfected cells; tRNA: riboprobe protected with yeast tRNA; M: marker; P: undigested probe.

In the first chimera, U3 sequence from HIV-1 NL4 were appended upstream of MLV transcribed sequences and 5’ ends of the intracellular RNAs this chimera produced were compared to those generated by MLV **(Fig. 3B)**. Strikingly, whereas MLV RNAs produced from the MLV promoter displayed a unique TSS, two MLV RNA isoforms were observed in cells transfected with the chimera containing HIV-1 U3 sequences upstream of MLV transcribed sequences **(Fig. 3B, lanes 2 and 3)**. Determining these RNAs’ 5’ ends revealed that their start sites’ positioning and heterogeneity matched what would be predicted if the major determinants of HIV-1 TSS heterogeneity were located upstream of the TSS, in U3 sequences **(Fig. 3A)**.

Two additional chimeras were generated to test whether MLV promoter sequences would generate unique TSS HIV-1 transcripts: one in which HIV-1 U3 sequences were replaced with MLV U3 at a spacing predicted to generate ^cap^3G RNAs exclusively, and a second designed to generate ^cap^1G RNAs if MLV U3 sequences dictated TSS multiplicity and positioning **(Fig. 3A)**. RNAs produced by these vectors were analyzed by CaDAL **(Fig. 3B)**. The results showed that HIV-1 RNAs transcribed from an MLV promoter each had a unique 5’ end **(Fig. 3B, lanes 5 and 6)**. Precise 5’ ends were confirmed by sequencing CaDAL products.

Virus-based competition experiments were then performed to address whether or not the MLV-driven unique 5’ end HIV-1 RNAs expressed from these latter chimeras performed the predicted functions **(Fig. 3C)**. Pairwise combinations of HIV-1 Ψ+ helper plasmid and test vectors (either the NL4-3-based vector used in **Fig. 1** or one of the two MLV/HIV-1 chimeras) were co-transfected into 293 T cells. RNA was prepared from both transfected cells and the viral particles they released, subjected to an RNase protection assay using a labeled probe that protected different length helper and vector fragments, and gel-separated products were analyzed by autoradiography **(Fig. 3C)**. The results indicated that as predicted, ^cap^1G RNAs were efficiently encapsidated but^cap^3G RNAs were excluded from packaging under these competitive conditions.

### HIV-1 heterogeneous TSSs do not result from tandem TATA-box use

As previously reported, the HIV-1 promoter contains a non-canonical TATA box with the sequence CATATAA [15]. This so-called CATA-box begins 29 bases upstream of the ^cap^3G RNA’s TSS and is highly conserved among HIV-1 strains. Other lentiviruses, including those studied here, differ from HIV-1 in possessing canonical TATA-box sequences **(Fig. 2A, 2B)**.

Experimental evidence confirming that CATA-box sequences (at positions -30 to -24) provide HIV-1’s TATA-box functions has been reported [15]. Nonetheless, because these sequences overlap from position -28 through -22 with sequences that contain a second TATA-like signal, it seemed conceivable that HIV-1’s heterogeneous TSS usage might reflect alternating use of two overlapping TATA elements **(Fig. 4A)**.

**Figure 4.**
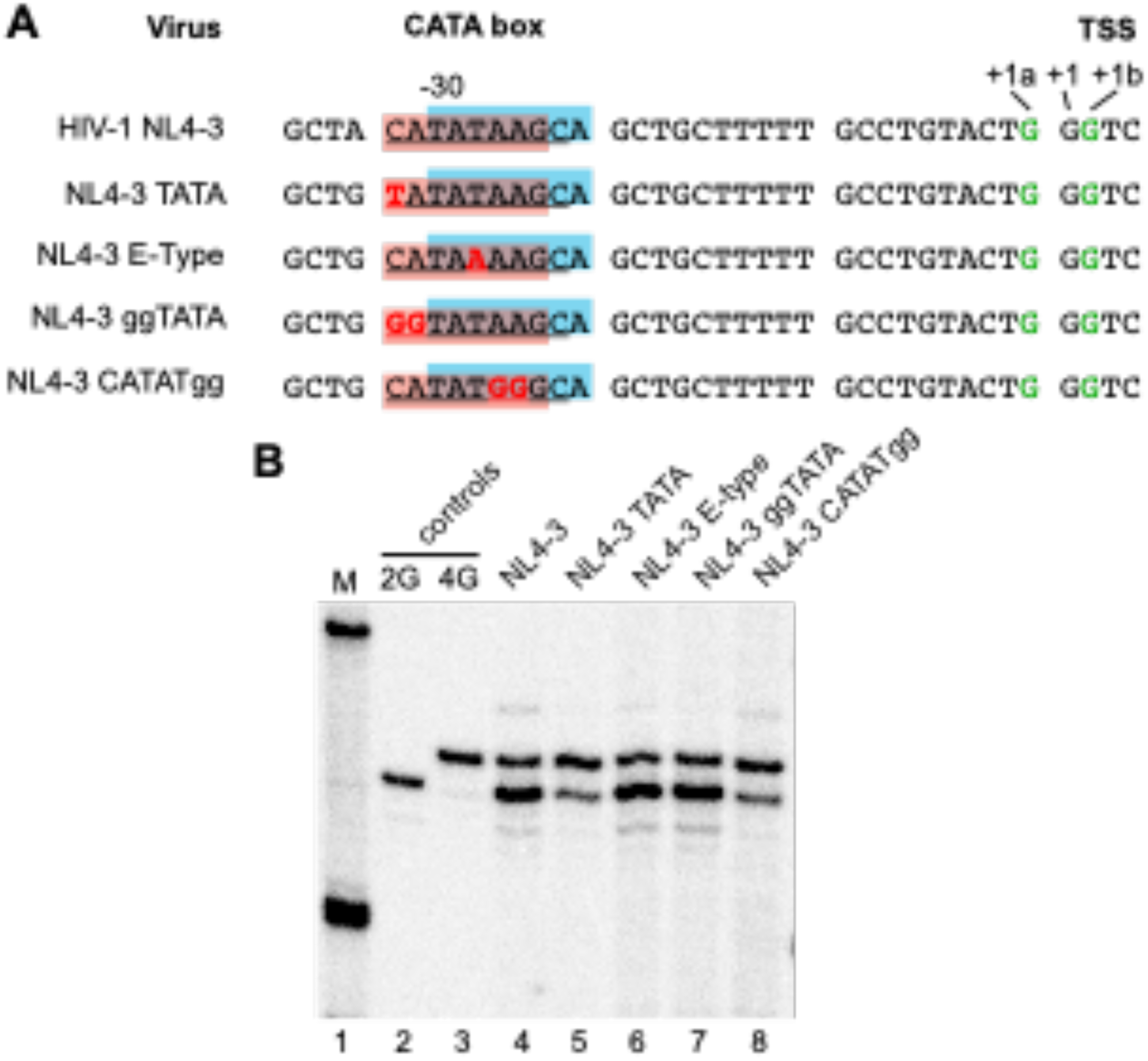
Testing tandem TATA-boxes roles in heterogeneous TSS usage (A) Mutations to test putative tandem TATA-boxes. Sequences from -35 to +3 of HIV-1 NL4-3 and mutated derivatives. The two potential TATA elements are shaded pink or blue. Mapped TSSs of NL4-3 and derivatives are in green. Mutated nucleotides are shown in red. (B) Single base resolution assay of HIV-1 cellular unspliced RNA 5’ ends assessed by CaDAL assay. M, size marker; lanes 2 and 3, products for 2G and 4G control plasmids; lanes 4-8, products of indicated cell RNA samples.

To examine contributions of these non-canonical core promoter elements to heterogeneous TSS usage, a point mutation was introduced to convert HIV-1’s CATA- box into a TATA-box **(Fig. 4A)**. 5’ end analysis revealed that TATA-NL4-3 continued to produce both ^cap^1G and ^cap^3G RNAs, albeit in a different ratio than the parental vector. Specifically, whereas about 70% of the parental vector’s RNAs were of the ^cap^1G form, about 55% of the TATA-box mutant’s RNAs had ^cap^3G ends **(Fig. 4B, lanes 4 and 5)**. Thus, single base conversion of the HIV-1 CATA-box into a canonical TATA-box did not ablate TSS usage heterogeneity but did modestly change RNA ratios.

To test the hypothesis that HIV-1’s CATA-box region functioned as a tandem TATA-box that caused heterogeneous TSS usage, mutations to ablate one or the other candidate TATA element were introduced. Notably, the third position in TATA-boxes is exceptionally conserved and exclusively occupied by T [15, 29, 30]. If the HIV-1 LTR had two overlapping TATA boxes, the third position of the second element would be at position -26. And indeed, sequences from -30 to -22 are highly conserved in all HIV-1 subtypes—including at the -26 position-- except in subtype E, which has a T/A polymorphism at position -26 [15, 31].

Consequently, if tandem TATA elements were responsible for heterogeneous transcription initiation, the A at -26 in subtype E viruses should render the second element inactive and subtype E viruses should produce only one RNA form. To test this notion, position -26 was changed to create an E subtype promoter **(Fig. 4A)**. 5’ end mapping showed that despite a potentially inactive second TATA-box, NL4-3 T-26A still produced two RNAs in a ratio similar to the parental vector **(Fig. 4B, lane 6)**.

Additional mutants were examined to further test the possible second TATA-like element. To inactivate the CATA-box while retaining the putative secondary TATA, CA residues at positions -30 and -29 were replaced with GG, creating NL4-3-GGTATA. To inactivate the potential second TATA-box, positions -25 and -24 were substituted with GG bases, creating NL4-3-CATATGG **(Fig. 4A)**. Examining these vectors RNAs’ revealed that both maintained heterogeneous TSS usage **(Fig. 4B)**. NL4-3 GGTATA produced ^cap^1G and ^cap^3G RNAs at levels similar to the parental vector, whereas NL4-3- CATATGG produced increased ^cap^3G RNAs **(Fig. 4B, lanes 7 and 8)**. Thus, whereas some CATA-box mutations affected RNA ratios, none disrupted heterogeneous TSS usage. and results were inconsistent with the hypothesis that HIV-1’s two TSSs reflect alternate use of two TATA elements.

### TSS-proximal sequences are key determinants of heterogeneous TSS usage

Because HIV-2 uses a unique TSS, a series of HIV-1 vectors with chimeric HIV-1/HIV-2 promoters were created to further map determinants of heterogeneous transcription initiation **(Fig.5A)**. Chimera 1 contained an intact HIV-2 U3 upstream of the HIV-1 GGG motif. Consistent with the findings above with MLV promoters upstream of HIV-1 sequences, placing the HIV-2 U3 promoter upstream of HIV-1 transcribed sequences resulted in the use of predominantly a single TSS, albeit with a small amount of residual secondary TSS use **(Fig. 5B)**.

**Figure 5.**
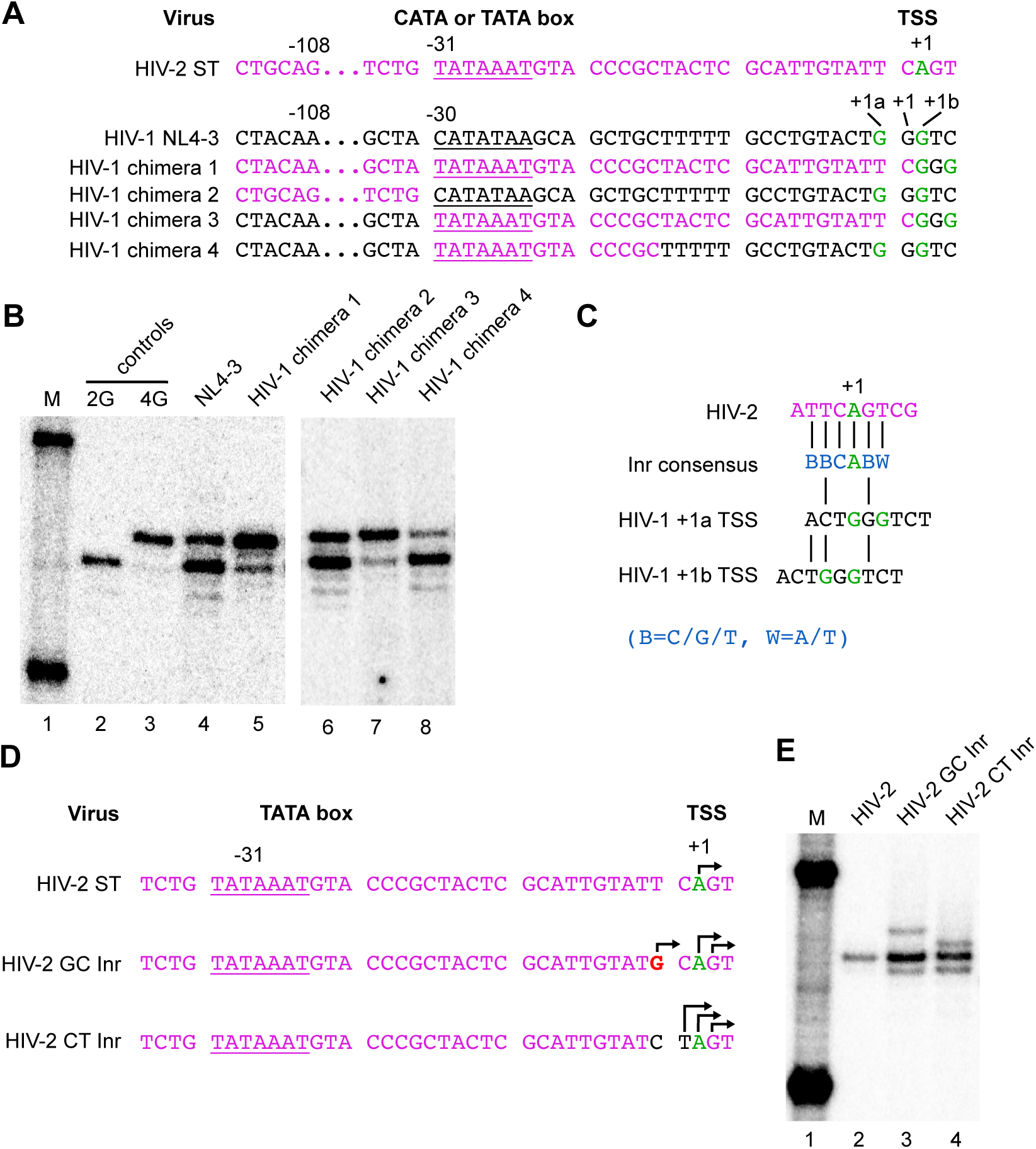
TSS usage in HIV-2/HIV-1 chimeric promoters and Inr variants. (A), Promoter sequences of NL4-3, HIV-2 ST and chimeric vectors. HIV-1 sequences are black and HIV-2 sequences are purple. CATA- or TATA-boxes are underlined. TSS in green. (B), Single base resolution assay of the cell RNA 5’ ends assessed by CaDEL. M, size marker; RNA samples from cells with indicated virus. (C) Alignment of HIV-2 and HIV-1 TSS sequences to Inr consensus. Inr consensus sequence [16] shown in blue, HIV-2 in purple, HIV-1 in black, TSS in green. HIV-1 sequence aligned to Inr consensus alternately using +1a as TSS or +1b as TSS. (D) Sequences of HIV-2 ST and Inr mutant core promoters. TATA-box underlined. Parental HIV-1 TSS in green. Observed TSS indicated with arrows. Insertion indicated by caret. Mutated nucleotides shown in red, or in black where derived from HIV-1. (E) Single base resolution assay of the cellular RNA 5’ ends assessed by CaDEL assay. M, size marker; RNA samples from cells with indicated viruses.

Additional chimeras were constructed to narrow down sequence elements important to multiple TSS use. Chimera 2 contained HIV-2 sequences upstream of the HIV-1 CATA- box followed by HIV-1 sequences, Chimera 3 had HIV-1 sequences upstream of HIV-2 TATA-box through -1 sequences, followed by HIV-1 transcribed sequences, and Chimera 4 was composed of HIV-1 sequences with a 15 bp fragment from HIV-2 replacing the CATA-box and adjacent region. Results with these chimeras **(Fig. 5C)** indicated that maintaining 15 bp of HIV-1 U3 sequences immediately adjacent to the TSS was sufficient to maintain dual TSS use, even when all upstream sequences were derived from HIV-2.

Previous work has described a conserved sequence element that flanks the TSSs of many Pol II promoters that display focal initiation. This motif is called the Initiator (Inr) and is believed to play a role in the precise positioning of Pol II at the TSS [8, 16]. Interestingly, sequences flanking the HIV-2 TSS are a perfect match to the Inr consensus, BBCA_+1_BW (**Fig.5D**). In contrast, sequence at the HIV-1 TSS bears no resemblance to the Inr consensus, although sequences in this region have been implicated as functionally important to HIV-1 transcriptional activation [32, 33].

To test whether HIV-2’s Inr was responsible for its unique TSS usage, two HIV-2 Inr region derivatives--both with substitutions based on HIV-1 sequences-- were designed and their transcripts’ 5’ ends determined (**Fig. 5E and F**). In one of these mutants, a T residue two bases upstream of the HIV-2 TSS was mutated to a G. This substitution is compatible with the canonical Inr sequence but introduced a purine as a potential secondary TSS at the -2 position, 29 bp downstream from the TATA-box. In the second mutant, the TC dinucleotide immediately upstream of the HIV-2 TSS were inverted to generate a CT, as is present in the corresponding position of HIV-1. This latter change is predicted to disrupt the Inr, because a C at position -1 is one of the most highly conserved Inr residues [16].

Examining intracellular RNAs revealed that both Inr region changes converted the HIV-2 U3 from a focal promoter to one that displayed heterogeneous TSSs (**Fig. 5F**). The -2 position G substitution maintained Inr homology but G_-2_ was used as a secondary TSS in addition to the parental A_+1_ start, and an unanticipated third TSS also was observed that mapped to G_+2_ (**Fig.9C, lane 3**). In the Inr disrupting TC inversion mutant (**Fig.9C, lane 4**), both A_+1_ and G_+2_ were used for transcription initiation, with a surprising third RNA isoform starting with a pyrimidine at T_-1_. These data showed that changes to the two nucleotides just upstream of the HIV-2 TSS, which introduced residues from analogous region of HIV-1, resulted in heterogeneous TSS usage. However, the role of these two nucleotides does not appear to reflect Inr functions, because both changes to this region that preserved HIV-2’s Inr consensus and those that ablated it displayed TSS heterogeneity.

### TSS proximal dinucleotide and core promoter element spacing focus TSS

The distance from the HIV-2 TATA-box to its +1 position is 31 bp, while the distance from the HIV-1 CATA box to +1a TSS is 29 bp and to +1b is 31 bp. Thus, published observations that reported TSSs shifted when the number of HIV-1 TSS G residues changed [1], paired with the fact that most eukaryotic Pol II transcripts begin with a purine [7, 9] raised the possibility that HIV-1 TSS choice was due primarily to core promoter element spacing relative to purine residues rather than to any specific properties of TSS-adjacent sequences.

To address this, promoter element spacing mutants were generated (**Fig. 6**). First, two dinucleotide insertion mutants were constructed that increased the distance between HIV-1’s CATA-box and its TSS (**Fig. 6A**). Both retained HIV-1’s purine-rich GGG trinucleotide motif and increased spacing to the CATA-box to 31 bp. However, one mutant was lengthened by a two bp duplication about 20 bp upstream of the TSS while the other was lengthened by an insertion of two residues from the corresponding region of HIV-2 immediately upstream of the GGG motif.

**Figure 6.**
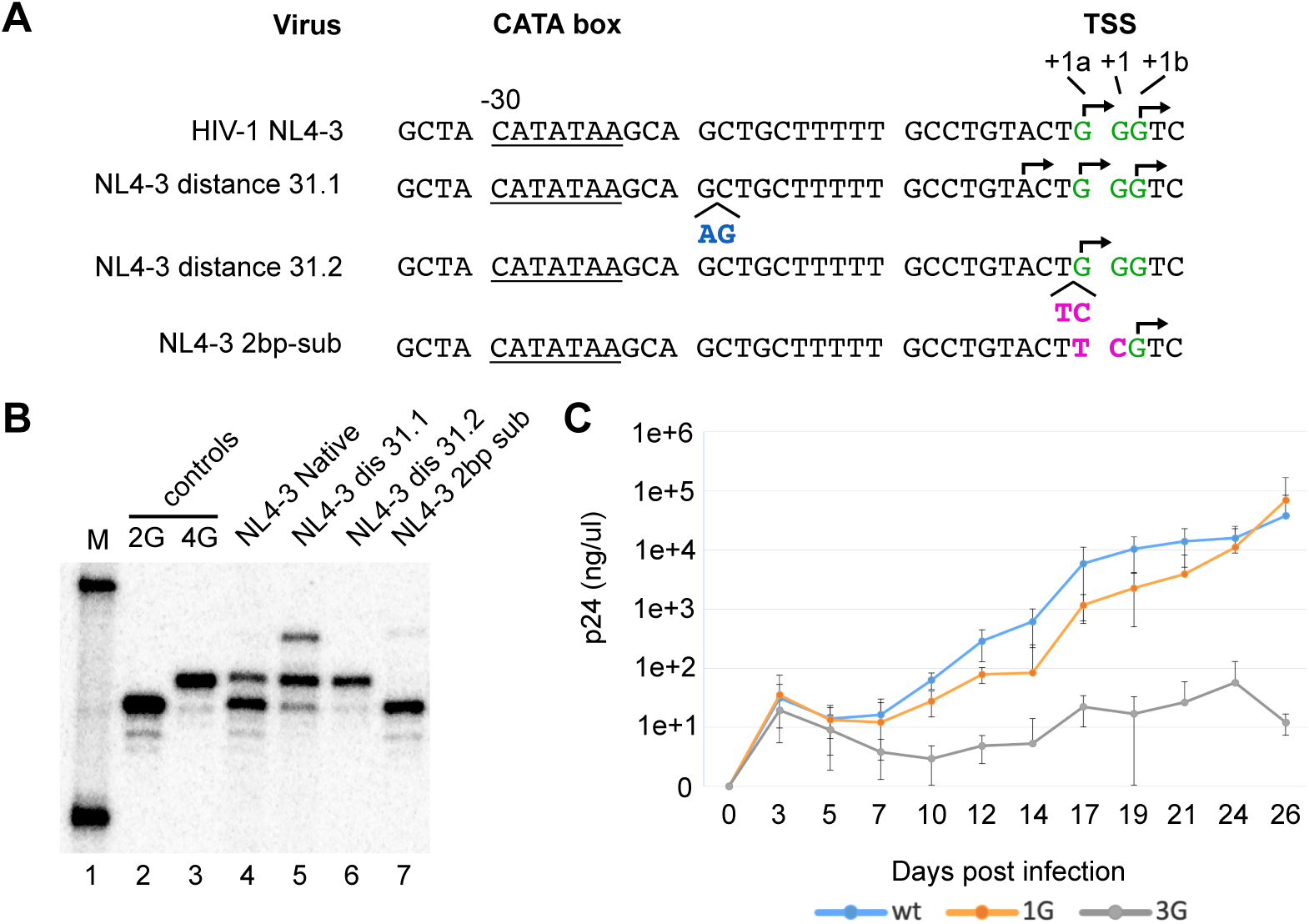
Influence of CATA-box distance and TSS-adjacent sequences on HIV-1 TSS selection. (A) Core promoter sequences of the wild-type HIV-1 NL4-3 and its mutants. CATA-box underlined. GGG trinucleotide motif indicated in green. Observed TSS indicated with arrows. Insertions indicated by caret. Inserted and mutated nucleotides shown in blue or in purple, where they correspond to nucleotides adjacent to TSS in HIV-2. (B), Single base resolution assay of the cellular RNA 5’ ends produced by mutated promoters, assessed by cap-dependent adaptor ligation/PCR method. M, molecular size marker; PCR products for 2G and 4G HIV-1 control plasmids are indicated; RNA samples extracted from cells transfected with corresponding virus are indicated. (C) Replication kinetics of parental NL4-3 and ^cap^1G RNAs-only and ^cap^3G RNAs-only variants. Virions were generated by transfection of 293T cells with proviral constructs and equal amounts (corresponding to 2 ng of CA p24) were used to infect CEM-SS cells. Virus production was monitored (see Material and Methods) at indicated time points. Data shown are from three independent infection experiments.

Determining TSS usage for these (**Fig.6B**) showed that the duplicated AG bases at positions -21 and -20 maintained heterogeneous TSS usage and added a third RNA start three bases upstream of +1a **(Fig.6B, lane 5).** Both TSSs of the parental NL4-3 vector were also still used, but the major RNA form shifted from ^cap^1G to ^cap^3G . These results suggested that nucleotide distance counting from the CATA-box contributed to specifying which residues served as HIV-1’s TSSs but was not the major determinant of heterogeneous TSS usage. In contrast, the promoter lengthened by the insertion of an HIV-2 derived TC dinucleotide immediately upstream of the GGG motif produced no ^cap^1G RNA. This later insertion converted the HIV-1 promoter into a focused promoter that almost exclusively produced ^cap^3G RNAs **(Fig. 6B, lane 6)**.

Based on these observations, an additional variant was constructed that reverted promoter element spacing to that of HIV-1 but maintained the HIV-2 derived dinucleotide as a two base substitution at HIV-1 positions +1a and +1 (**Fig. 6A** and **6B lane 7**). Results with this variant indicated that introducing two substitutions into HIV-1’s TSS region were sufficient to convert HIV-1 from heterogeneous TSS use to initiating transcription at a unique position.

These final two mutations—one a two-base insertion, and the second a two-base substitution—were thus each sufficient to convert HIV-1 from a heterogeneous TSS using virus to a unique TSS using virus: the first expressing only ^cap^3G RNAs and the second only ^cap^1G RNAs. To test their replication properties, these two mutant promoter regions were incorporated into both LTRs of a fully infectious HIV-1 NL4-3 molecular clone. Virions were produced by transfecting 293T cells, normalized by RT levels, and used to infect CEM-SS cells, The results of monitoring virus spread over time (**Fig. 6C**) indicated that the ^cap^1G-only virus had delayed replication kinetics compared to parental NL4-3 virus (Fig. 6C), although replication recovered and virus production reached parental NL4-3 levels after four weeks of passage. In contrast, the ^cap^3G-only mutant virus showed significant replication deficiencies. Its very low virus production persisted throughout the four week experiment (Fig.6C).

## Discussion

HIV-1 subtypes A and B have previously been shown to initiate transcription at two distinct template positions and generate two major RNA isoforms: ^cap^1G and ^cap^3G [1, 2]. These RNAs adopt distinct folded structures that differ in their abilities to support specific replication processes, and thus heterogeneous TSS usage regulates HIV-1 RNA fates. The two HIV-1 TSSs consist of the first and the third residues in a GGG motif located in the viral LTR, with the ^cap^1G RNA form serving as virion genomic RNA and ^cap^3G RNAs translated to yield viral proteins. Due to its GGG trinucleotide motif and two TSSs separated by one nucleotide, the HIV-1 promoter resembles the so-called twin-TSS promoters first characterized from a high-throughput screen of mammalian transcripts [9].

Here, database analysis showed that the TSS motif is highly conserved among HIV-1 family members. This prompted an experimental analysis of TSS usage in additional HIV-1 strains and a broader range of retroviruses. The results showed that all tested HIV-1 strains, including several different subtypes and a group O isolate, produced two RNA forms in cells, only one of which was observed in viral particles. Thus, the control of RNA fates by heterogeneous TSS usage that has been described for subtype A and B isolates appears to be a conserved feature of HIV-1 viruses.

In contrast, all other retroviruses tested displayed only a single 5’ end isoform, both in cells and in virions. This was true for the simple retroviruses RSV, RD114, and MLV as well as for the lentiviruses HIV-2, SIVmac, and FIV. RSV and MLV have previously been found to use single TSSs based on virion RNA sequence analysis [26, 27]. Another report described some heterogeneity at the 5’ end of MLV RNA extracted from viral particles or transcribed in cell lysates, but these were assayed by S1 nuclease protection and the authors concluded that there may have been partial DNA probe protection by the RNA 5’ cap [34]. Pol II transcription generally initiates at purine residues [7, 9], and because some of the viruses tested here have polypurine runs in their TSS regions (GGGG in RD114, or GAGGAG in FIV) it seemed possible that they might employ more than one TSS. However, the cap-dependent adapter ligation/PCR assay used here revealed that each of these used a unique TSS, and that the same single RNA form was observed in both cells and virions.

Thus, TSS heterogeneity is not an obligatory feature of retroviral replication but instead one that distinguishes HIV-1 from all other tested retroviruses, suggesting that other retroviruses must use strategies different from HIV-1’s to control their unspliced RNAs’ functions. As a possible example of this, recently it was shown that the cytoplasmic fates of full-length MLV RNA could be determined by which nuclear export pathway and nuclear export factors were recruited to viral RNA [35]. Specifically, it has been reported that binding of NXF1 and SRp20 to full-length MLV RNA drives the RNA through NXF1 dependent nuclear export to polysomes [36], whereas interaction with CRM1 drives viral RNA to virion assembly sites through CRM1 dependent nuclear export [35].

Interestingly, while this article was in preparation, another group examined TSS usage in lentiviruses [37]. That paper confirmed heterogeneous TSS usage in multiple HIV-1 strains but differed from the current study in its HIV-2 conclusions, which indicated that HIV-2 displays heterogeneous TSSs [37]. We have not determined the cause of this discrepancy but note that while HIV-2 appeared to use a single TSS here, we did observe some TSS heterogeneity when HIV-1 sequences were expressed using the HIV-2 U3 promoter (Chimera 1 in **Fig. 5**). Additionally, our two reports employed different technologies to map RNA 5’ ends. To reduce artifacts and enhance specificity, researchers who perform genome-wide analyses of transcription start sites have now largely replaced the 5’ RACE approach used in the other study with CAGE (cap analysis gene expression) or other cap-dependent approaches like those used in our study [38].

Here, the determinants of HIV-1 heterogeneous TSS usage were mapped using targeted mutations and promoter chimeras. Previous work had shown that introducing mutations into the HIV-1 GGG motif did not affect heterogeneous TSS usage [1], and work here confirmed that the naturally arising TSS variant AGG also exhibited heterogeneous TSS usage. Thus, because determinants of TSS use appeared not to reside in the start sites themselves, promoter swaps were constructed to test if the determinants were located upstream or downstream of HIV-1’s GGG motif. Analyzing RNA 5’ ends revealed that when MLV transcripts were produced using an HIV-1 promoter, two major TSSs separated by one base were employed. In contrast, HIV-1 RNAs produced using the MLV U3 promoter displayed only one TSS, independent of the number of TSS proximal purines.

Having mapped determinants to upstream of the TSS, additional chimeras were generated to test the influence of specific promoter elements. The best characterized core promoter element is the TATA-box, with a consensus sequence of TATAWAAR that is located between -33/-26 to -27/-18 upstream of the TSS [7, 9, 13]. Many Pol II promoters, including many that support single or twin TSS use, do not have TATA-boxes [8, 9]. HIV-1 contains a non-canonical TATA-box with the sequence CATATAA, which has been dubbed a CATA-box [15]. CATA-box sequences are highly conserved among HIV-1s and found in a limited number of human promoters, including those for β-globin and IL1B [14]. CATA-boxes are associated with lower levels of transcriptional activity [29, 30], lower TBP binding affinity [14, 29], and less stable TFIIA-TBP-DNA complexes [39, 40]. Previously it was shown that converting the CATA-box into a canonical TATA-box led to elevated levels of HIV-1 promoter expression and enhanced fitness of the virus in chronically infected cell culture [15].

Examining the sequence of HIV-1’s CATA-box suggested the possibility that two overlapping TATA-like elements in the HIV-1 core promoter might be responsible for dual TSS use. However, a series of CATA-box region mutants revealed no evidence for the use of two TATA-like elements, although some CATA-box mutants altered transcript isoform ratios.

Additional work indicated that spacing between the CATA/TATA-box and TSS is an important, but not the only, factor in defining lentiviral TSSs. Start site selection through distance counting is characteristic of TATA-box containing promoters [9, 41]. In HIV-1, the +1a and +1b TSSs are located 29 and 31 bp from the CATA-box, respectively. Here, one 2 bp insertion between the CATA-box and TSS changed the ratios of HIV-1 ^cap^1G and ^cap^3G RNAs and lead to the appearance of a third RNA that initiated in the first purine upstream of original TSS. However, whereas observations with this mutant suggested that distance counting specified start site selection, this mechanism appeared to be over-ridden in a different mutant, in which the CATA-TSS distance was lengthened by a two nucleotide insertion directly upstream of the GGG motif. Observations with this later mutant suggested that TSS-proximal residues play a dominant role in determining whether or not TSS heterogeneity is observed.

These TSS-proximal sequences reside in the same location as an element involved in some promoters’ TSS selection. The so-called Initiator (Inr) element overlaps the TSS and may function in the precise positioning of Pol II at the TSS [8, 16]. Most focused promoters regulated by Inr do not contain TATA-boxes, although a significant proportion contain both a TATA-box and Inr [8, 16]. Promoters containing both TATA-box and Inr elements differ from those containing only one of these elements in terms of promoter strength [42] and responses to transcription factors such as NC2 [43] and HMGA1 [44]. HIV-1 TSS-flanking sequences, mapped alternately to positions -6-+8 [32] or -6-+11 [33], are important to transcriptional activation, but HIV-1 lacks the BBCA_+1_BW Inr consensus sequence [16]. In contrast, HIV-2’s single TSS is flanked by a perfect match to the Inr consensus.

To test if the presence or absence of an Inr dictated TSS heterogeneity, HIV-1 / HIV-2 core promoter chimeras were tested. Because one mutation that disrupted HIV-2’s perfect Inr consensus converted it from a focused promoter into one that displayed heterogeneous starts, initial results suggested that the presence or absence of a canonical Inr element might dictate TSS heterogeneity. However, subsequent experimentation showed that whereas determinants reside in the same genetic interval as Inr elements, the regulation of lentivirus TSS heterogeneity appears mechanistically distinct from Inr-mediated regulation.

The observations here that all retroviruses other than HIV-1 use unique transcription start sites suggest that HIV-1’s dependence on two TSSs is a relatively recent acquisition. Results of the promoter element spacing experiments above suggested that the ^cap^1G TSS is HIV-1’s ancestral TSS and that the TSS that generates ^cap^3G RNA is a newer acquisition. This notion is consistent with experimental observations that ^cap^1G RNAs can function both in genomic RNA packaging and as mRNAs, whereas ^cap^3G RNAs are limited to intracellular roles and excluded from packaging [1, 3]. Similarly, the initial infectivity studies performed here show that neither variant that expresses only one HIV-1 RNA is fully infectious but that replication of the ^cap^3G-only variant is the more severely impaired. Additional work will be required to determine how these two RNAs and the functions they support act in concert to support HIV-1 replication.

## Methods

### Sequences alignment and evolutionary analysis by Maximum Likelihood

Conservation of TSS sequence among SIVcpz/HIV-1 viruses was addressed using Web Alignment and Consensus Maker tools at Los Alamos HIV-1 Sequences Database (https://www.hiv.lanl.gov/content/index). Alignment was performed for positions 450-460 (HXB2 reference sequence coordinates) of the 5’ LTR (573 available sequences) or 9536-9546 of 3’ LTR (2777 available sequences). Consensus sequence logo was generated by alignment of 573 available 5’ LTR sequences in WebLogo software [45].

Sequence alignment and reconstruction of phylogenetic relationships was performed using MEGA v.10 software [46]. ClustalW was used for alignment and the evolutionary history was inferred using Maximum Likelihood and a General Time Reversible model [47] with gamma distribution. Confidence of phylogenetic tree was determined through 1,000 bootstrap permutations and included FIV as an outgroup.

### Plasmids

Plasmids containing molecular clones of the following were obtained from the NIH HIV Reagent Program: HIV-1 subtype C (pMJ4); SIVmac239-ΔNef and HIV-2 ST. “Native” refers to a previously described NL4-3 based vector that contains LTRs, 688 bp of HIV- 1 5’ leader, RRE, and a puromycin resistance cassette [48]. DNA fragments containing 5’ LTR and leader sequences through approximately 250bp of *gag* for the following viruses: HIV-1 group M subtype D; HIV-1 group O; CRF60_BC; SIVcpzEK505 and SIVcpzMT145 (accession numbers AB485649; KU168298; KC899079; DQ373065 and JN835462 correspondently) synthesized by GENEWIZ and cloned into Native in place of NL4-3 sequences, to generate D-Native, O-Native, CRF60-Native, EK-Native and MT-Native. CMVΔR8.2 is served as the Ψ- helper [49]. The Ψ+ helper, aka NL4-3 GPP, replication-defective HIV-1 NL4-3 with deleted *vpu*, *vpr*, *nef* and partial deletion of *env* was described previously [4]. pNL4-3 contains an infectious NL4-3 strain HIV-1 molecular clone [50]. Infectious RSV molecular clone pRC.V8 was kindly provided by Leslie Parent; its *src* gene is replaced with a hygromycin resistance gene.

Fragments containing targeted mutations were generated using overlap extension PCR.

### Cells, viruses, transfections, and infections

174xCEM and CRFK FIV AZT-1 cells were obtained from the NIH HIV Reagent Program. 293T, CEM-SS and DF1 cells were purchased from ATCC. To produce HIV-1, HIV-2 and SIV viruses, freshly seeded 293T cells grown in DMEM supplemented with 10% FBS + 50 µg/mL gentamycin at 37°C with 5% CO2 (hereafter, supplemented DMEM) were transfected using polyethylenimine (Polysciences) [51]. For infectious molecular clones 5 µg of plasmid DNA were used for transfection, while Native and derivatives were co-transfected with CMVΔR8.2 at a 3:1 molar ratio (8 µg of plasmid DNA total). Note that HIV-2 based vectors were mobilized using an HIV-1 helper since HIV-1 Tat can efficiently activate transcription from the HIV-2 promoter [52, 53].

To establish cells chronically infected with SIVmac239-ΔNef, 174xCEM cells were grown in RPMI 1640 supplemented with 10% FBS + 50 µg/mL gentamycin at 37°C with 5% CO2. Media, containing approximately 2 ng CA-p24 of SIVmac239-ΔNef virus was added to 4 × 10^6^ 174xCEM cells. Infection was monitored by quantitative PCR-based reverse transcriptase (RT) assay, in which HIV-1–containing medium with known concentrations of CA-p24 was used as the standard [54]. Virus and cells were harvested after virus levels plateaued (approx. 12 days).

To produce FIV, cat kidney cells chronically infected with FIV (CRFK FIV AZT-1) were grown in a 1:1 mixture of Liebovitz (L-15) medium and supplemented DMEM in the presence of 10 µg/mL AZT. After cells reached confluency, virus and cells were harvested.

CRFK cells constitutively produce infectious RD-114 virus [55]. To establish a 293T culture chronically infected with RD-114, CRFK FIV AZT-1 cells were grown without AZT. Viral media containing 8 µg/mL polybrene was applied to 293T cells at 50% confluency. After 6 hours, media was replaced with fresh supplemented DMEM and cells were passaged for two weeks. Infection was monitored by RT assay [54]. After virus levels plateaued, virus and cells were harvested.

NIH3T3 cells chronically infected with MLV [19] were maintained in supplemented DMEM. After cells reached confluency, virus and cells were harvested.

To establish DF1 cell (immortalized chicken embryo fibroblasts) chronically infected with RSV, DF1 cells were grown in supplemented DMEM plus 1mM sodium pyruvate. DF1cells were transfected with 5 µg pRC.V8 as described previously [56]. 48 hours after transfection, filtered viral media was added to fresh 50% confluent DF1 cells, and cells were passaged. Virus production monitored by RT assay. After virus levels plateaued, virus and cells were harvested.

For HIV-1 virus growth assay pNL4-3 and its mutated versions pNL4-3 TC-1G and pNL4-3 TC-3G were transfected into freshly seeded 293T cells using polyethylenimine (Polysciences) [51]. Virus containing media were harvested 48 hours post transfection and passed through 0.22-μm filters. Virus level in harvested media was quantified by RT assay and virus equivalent to 2 ng CA-p24 was used to infect 2 × 10^6^ CEM-SS cells grown in RPMI 1640 media supplemented with 10% fetal bovine serum and 50 µg/mL gentamicin. Infected CEM-SS cells were passed for approximately 5 weeks. Monitoring the level of virus production was performed by RT assay in the media aliquots harvested every 2^nd^ or 3^rd^ day.

### RNA extraction, CaDAL assay, Sanger sequencing, RNAse protection assay

Virus particles were collected by centrifuging filtered culture media through a 20% sucrose cushion at 25,000 rpm for 2 hrs. Virus pellet and cell RNA was isolated using TRIzol according to the manufacturer’s protocol (Ambion) and DNase treated. 5′ ends of capped viral RNAs were analyzed using a **ca**p-**d**epended **a**daptor **l**igation/PCR (CaDAL) assay [3, 24]. Briefly, RNA extracted from viral particles or virus producing cells was used as a template for cDNA synthesis using the TeloPrime Full-Length cDNA Amplification Kit V2 (Lexogen) according to the manufacturer’s protocol. Each cDNA preparation was primed with virus-specific primers complementary to a region of viral RNA downstream of its major 5’ splice site (Table 1). Cap-depended adaptor ligation and second-strand synthesis was performed according to the manufacturer’s instructions. Viral cDNA was amplified for 15 cycles with Phusion polymerase (NEB) using a primer complementary to the adaptor (TGGATTGATATGTAATACGACTCACTATAGG) and one complementary to the R region of each virus (Table 1). Prior to PCR, the adaptor complementary primer was labeled with [γ-32P]-ATP and T4 PNK (NEB) using the manufacturer’s protocol. PCR products were visualized by phosphorimaging after resolution by denaturing 15% PAGE at 50W for ∼5 h. For Sanger sequencing, the same products were amplified using non-radiolabeled primers, extracted from agarose using the NEB Monarch DNA gel extraction kit, and cloned using a TOPO TA kit (Thermofisher). At least 12 individual clones were sequenced for each virus sample.

**Table 1.**
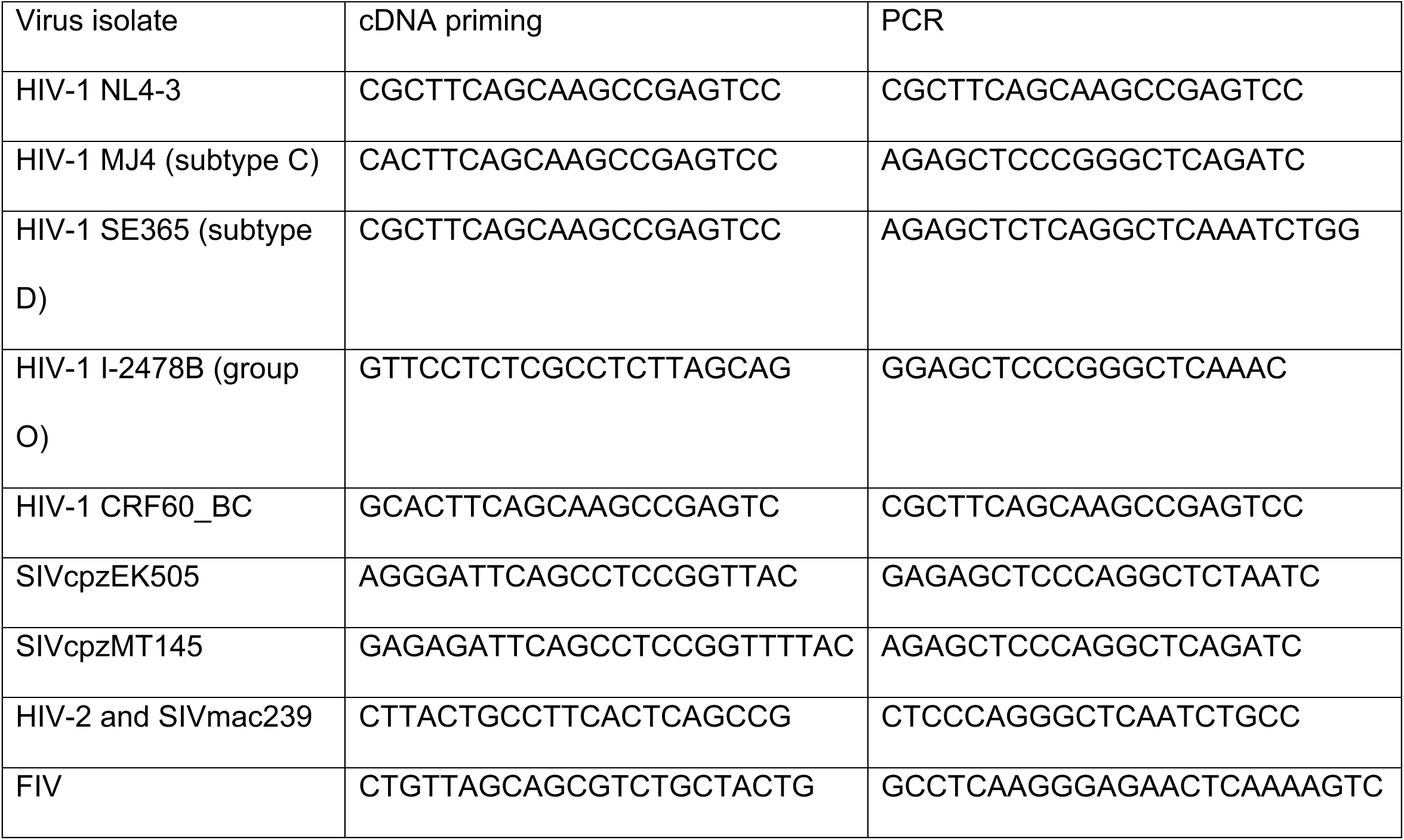
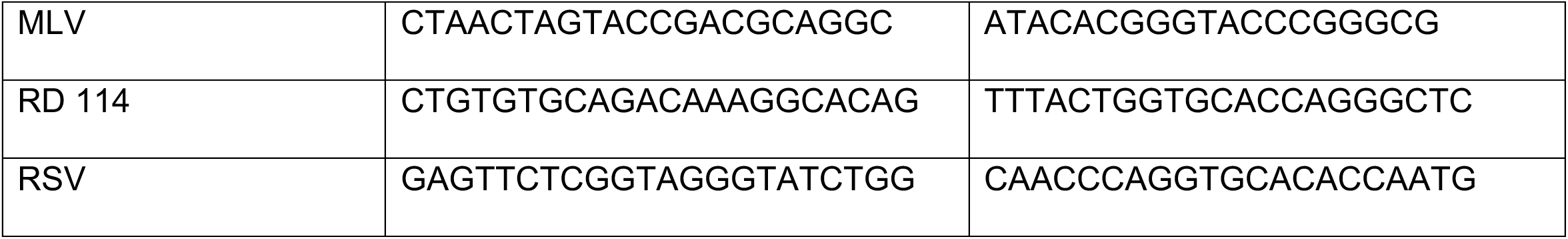
Virus specific primers used in cap-depended ligation/PCR protocol.

RNase Protection assay (RPA) was performed as previously described [6]. The riboprobe HIVgag/CMV used in this study was described previously [6] and targets a 201-nt fragment of *gag* unique to the NL4-3 GPP helper and 289 nt of CMV promoter sequence from *puro*-expressing cassette unique to test vectors.

## Acknowledgments

We thank Nastassia Vlasava for help with phylogenetic analysis. The research was supported by NIH grants R01 AI50498 and U54 AI170660 to AT.

## Author Contributions

SK Designed and performed experiments and wrote the manuscript

CB and KG Performed experiments and contributed to manuscript preparation

AT Designed experiments and wrote the manuscript

## References

[1] Kharytonchyk S, Monti S, Smaldino PJ, Van V, Bolden NC, Brown JD, et al. Transcriptional start site heterogeneity modulates the structure and function of the HIV- 1 genome. Proc Natl Acad Sci U S A. 2016;113:13378–83.

[2] Masuda T, Sato Y, Huang YL, Koi S, Takahata T, Hasegawa A, et al. Fate of HIV-1 cDNA intermediates during reverse transcription is dictated by transcription initiation site of virus genomic RNA. Sci Rep. 2015;5:17680.

[3] Esquiaqui JM, Kharytonchyk S, Drucker D, Telesnitsky A. HIV-1 spliced RNAs display transcription start site bias. RNA. 2020;26:708–14.

[4] Lu K, Heng X, Garyu L, Monti S, Garcia EL, Kharytonchyk S, et al. NMR detection of structures in the HIV-1 5’-leader RNA that regulate genome packaging. Science. 2011;334:242–5.

[5] Brown JD, Kharytonchyk S, Chaudry I, Iyer AS, Carter H, Becker G, et al. Structural basis for transcriptional start site control of HIV-1 RNA fate. Science. 2020;368:413–7.

[6] Ding P, Kharytonchyk S, Kuo N, Cannistraci E, Flores H, Chaudhary R, et al. 5’-Cap sequestration is an essential determinant of HIV-1 genome packaging. Proc Natl Acad Sci U S A. 2021;118.

[7] Carninci P, Sandelin A, Lenhard B, Katayama S, Shimokawa K, Ponjavic J, et al. Genome-wide analysis of mammalian promoter architecture and evolution. Nat Genet. 2006;38:626–35.

[8] Kadonaga JT. Perspectives on the RNA polymerase II core promoter. Wiley Interdiscip Rev Dev Biol. 2012;1:40–51.

[9] Ponjavic J, Lenhard B, Kai C, Kawai J, Carninci P, Hayashizaki Y, et al. Transcriptional and structural impact of TATA-initiation site spacing in mammalian core promoters. Genome Biol. 2006;7:R78.

[10] Andersson R, Sandelin A. Determinants of enhancer and promoter activities of regulatory elements. Nat Rev Genet. 2020;21:71–87.

[11] Haberle V, Stark A. Eukaryotic core promoters and the functional basis of transcription initiation. Nat Rev Mol Cell Biol. 2018;19:621–37.

[12] Danino YM, Even D, Ideses D, Juven-Gershon T. The core promoter: At the heart of gene expression. Biochim Biophys Acta. 2015;1849:1116–31.

[13] Juven-Gershon T, Hsu JY, Theisen JW, Kadonaga JT. The RNA polymerase II core promoter - the gateway to transcription. Curr Opin Cell Biol. 2008;20:253–9.

[14] Savinkova L, Drachkova I, Arshinova T, Ponomarenko P, Ponomarenko M, Kolchanov N. An experimental verification of the predicted effects of promoter TATA-box polymorphisms associated with human diseases on interactions between the TATA boxes and TATA-binding protein. PLoS One. 2013;8:e54626.

[15] van Opijnen T, Kamoschinski J, Jeeninga RE, Berkhout B. The human immunodeficiency virus type 1 promoter contains a CATA box instead of a TATA box for optimal transcription and replication. J Virol. 2004;78:6883–90.

[16] Vo Ngoc L, Cassidy CJ, Huang CY, Duttke SH, Kadonaga JT. The human initiator is a distinct and abundant element that is precisely positioned in focused core promoters. Genes Dev. 2017;31:6–11.

[17] Berkhout B, Vastenhouw NL, Klasens BI, Huthoff H. Structural features in the HIV-1 repeat region facilitate strand transfer during reverse transcription. RNA. 2001;7:1097–114.

[18] Ohi Y, Clever JL. Sequences in the 5’ and 3’ R elements of human immunodeficiency virus type 1 critical for efficient reverse transcription. J Virol. 2000;74:8324–34.

[19] Kulpa D, Topping R, Telesnitsky A. Determination of the site of first strand transfer during Moloney murine leukemia virus reverse transcription and identification of strand transfer-associated reverse transcriptase errors. EMBO J. 1997;16:856–65.

[20] Topping R, Demoitie MA, Shin NH, Telesnitsky A. Cis-acting elements required for strong stop acceptor template selection during Moloney murine leukemia virus reverse transcription. J Mol Biol. 1998;281:1–15.

[21] Hemelaar J. The origin and diversity of the HIV-1 pandemic. Trends Mol Med. 2012;18:182–92.

[22] Yamaguchi J, Vallari A, McArthur C, Sthreshley L, Cloherty GA, Berg MG, et al. Brief Report: Complete Genome Sequence of CG-0018a-01 Establishes HIV-1 Subtype L. J Acquir Immune Defic Syndr. 2020;83:319–22.

[23] Keele BF, Van Heuverswyn F, Li Y, Bailes E, Takehisa J, Santiago ML, et al. Chimpanzee reservoirs of pandemic and nonpandemic HIV-1. Science. 2006;313:523–6.

[24] Ohtake H, Ohtoko K, Ishimaru Y, Kato S. Determination of the capped site sequence of mRNA based on the detection of cap-dependent nucleotide addition using an anchor ligation method. DNA Res. 2004;11:305–9.

[25] McAllister RM, Nicolson M, Gardner MB, Rongey RW, Rasheed S, Sarma PS, et al. C-type virus released from cultured human rhabdomyosarcoma cells. Nat New Biol. 1972;235:3–6.

[26] Rose JK, Haseltine WA, Baltimore D. 5’-terminus of Moloney murine leukemia virus 35s RNA is m7G5’ ppp5’ GmpCp. J Virol. 1976;20:324–9.

[27] Keith J, Fraenkel-Conrat H. Identification of the 5’ end of Rous sarcoma virus RNA. Proc Natl Acad Sci U S A. 1975;72:3347–50.

[28] Apetrei C, Robertson DL, Marx PA. The history of SIVS and AIDS: epidemiology, phylogeny and biology of isolates from naturally SIV infected non-human primates (NHP) in Africa. Front Biosci. 2004;9:225–54.

[29] Hoopes BC, LeBlanc JF, Hawley DK. Contributions of the TATA box sequence to rate-limiting steps in transcription initiation by RNA polymerase II. J Mol Biol. 1998;277:1015–31.

[30] Wobbe CR, Struhl K. Yeast and human TATA-binding proteins have nearly identical DNA sequence requirements for transcription in vitro. Mol Cell Biol. 1990;10:3859–67.

[31] De Baar MP, De Ronde A, Berkhout B, Cornelissen M, Van Der Horn KH, Van Der Schoot AM, et al. Subtype-specific sequence variation of the HIV type 1 long terminal repeat and primer-binding site. AIDS Res Hum Retroviruses. 2000;16:499–504.

[32] Jones KA, Luciw PA, Duchange N. Structural arrangements of transcription control domains within the 5’-untranslated leader regions of the HIV-1 and HIV-2 promoters. Genes Dev. 1988;2:1101–14.

[33] Zenzie-Gregory B, Sheridan P, Jones KA, Smale ST. HIV-1 core promoter lacks a simple initiator element but contains a bipartite activator at the transcription start site. J Biol Chem. 1993;268:15823–32.

[34] Fuhrman SA, Van Beveren C, Verma IM. Identification of a RNA polymerase II initiation site in the long terminal repeat of Moloney murine leukemia viral DNA. Proc Natl Acad Sci U S A. 1981;78:5411–5.

[35] Mougel M, Akkawi C, Chamontin C, Feuillard J, Pessel-Vivares L, Socol M, et al. NXF1 and CRM1 nuclear export pathways orchestrate nuclear export, translation and packaging of murine leukaemia retrovirus unspliced RNA. RNA Biol. 2020;17:528–38.

[36] Bartels H, Luban J. Gammaretroviral pol sequences act in cis to direct polysome loading and NXF1/NXT-dependent protein production by gag-encoded RNA. Retrovirology. 2014;11:73.

[37] Rawson JMO, Nikolaitchik OA, Shakya S, Keele BF, Pathak VK, Hu WS. Transcription Start Site Heterogeneity and Preferential Packaging of Specific Full-Length RNA Species Are Conserved Features of Primate Lentiviruses. Microbiol Spectr. 2022;10:e0105322.

[38] Adiconis X, Haber AL, Simmons SK, Levy Moonshine A, Ji Z, Busby MA, et al. Comprehensive comparative analysis of 5’-end RNA-sequencing methods. Nat Methods. 2018;15:505–11.

[39] Stewart JJ, Fischbeck JA, Chen X, Stargell LA. Non-optimal TATA elements exhibit diverse mechanistic consequences. J Biol Chem. 2006;281:22665–73.

[40] Stewart JJ, Stargell LA. The stability of the TFIIA-TBP-DNA complex is dependent on the sequence of the TATAAA element. J Biol Chem. 2001;276:30078–84.

[41] Smale ST, Kadonaga JT. The RNA polymerase II core promoter. Annu Rev Biochem. 2003;72:449–79.

[42] Lo K, Smale ST. Generality of a functional initiator consensus sequence. Gene. 1996;182:13–22.

[43] Malecova B, Gross P, Boyer-Guittaut M, Yavuz S, Oelgeschlager T. The initiator core promoter element antagonizes repression of TATA-directed transcription by negative cofactor NC2. J Biol Chem. 2007;282:24767–76.

[44] Xu M, Sharma P, Pan S, Malik S, Roeder RG, Martinez E. Core promoter-selective function of HMGA1 and Mediator in Initiator-dependent transcription. Genes Dev. 2011;25:2513–24.

[45] Crooks GE, Hon G, Chandonia JM, Brenner SE. WebLogo: a sequence logo generator. Genome Res. 2004;14:1188–90.

[46] Kumar S, Stecher G, Li M, Knyaz C, Tamura K. MEGA X: Molecular Evolutionary Genetics Analysis across Computing Platforms. Mol Biol Evol. 2018;35:1547–9.

[47] Nei M, Kumar S. Molecular evolution and phylogenetics. Oxford ; New York: Oxford University Press; 2000.

[48] Heng X, Kharytonchyk S, Garcia EL, Lu K, Divakaruni SS, LaCotti C, et al. Identification of a minimal region of the HIV-1 5’-leader required for RNA dimerization, NC binding, and packaging. J Mol Biol. 2012;417:224–39.

[49] Naldini L, Blomer U, Gallay P, Ory D, Mulligan R, Gage FH, et al. In vivo gene delivery and stable transduction of nondividing cells by a lentiviral vector. Science. 1996;272:263–7.

[50] Adachi A, Gendelman HE, Koenig S, Folks T, Willey R, Rabson A, et al. Production of acquired immunodeficiency syndrome-associated retrovirus in human and nonhuman cells transfected with an infectious molecular clone. J Virol. 1986;59:284–91.

[51] Boussif O, Lezoualc’h F, Zanta MA, Mergny MD, Scherman D, Demeneix B, et al. A versatile vector for gene and oligonucleotide transfer into cells in culture and in vivo: polyethylenimine. Proc Natl Acad Sci U S A. 1995;92:7297–301.

[52] Emerman M, Guyader M, Montagnier L, Baltimore D, Muesing MA. The specificity of the human immunodeficiency virus type 2 transactivator is different from that of human immunodeficiency virus type 1. EMBO J. 1987;6:3755–60.

[53] Fenrick R, Malim MH, Hauber J, Le SY, Maizel J, Cullen BR. Functional analysis of the Tat trans activator of human immunodeficiency virus type 2. J Virol. 1989;63:5006–12.

[54] Vermeire J, Naessens E, Vanderstraeten H, Landi A, Iannucci V, Van Nuffel A, et al. Quantification of reverse transcriptase activity by real-time PCR as a fast and accurate method for titration of HIV, lenti- and retroviral vectors. PLoS One. 2012;7:e50859.

[55] Yoshikawa R, Sato E, Igarashi T, Miyazawa T. Characterization of RD-114 virus isolated from a commercial canine vaccine manufactured using CRFK cells. J Clin Microbiol. 2010;48:3366–9.

[56] Pfeiffer JK, Topping RS, Shin NH, Telesnitsky A. Altering the intracellular environment increases the frequency of tandem repeat deletion during Moloney murine leukemia virus reverse transcription. J Virol. 1999;73:8441–7.

